# ERβ limits T cell-mediated inflammation to maintain immune homeostasis

**DOI:** 10.1101/2025.06.21.660869

**Authors:** Sarah K. McNeer, Alyssia V. Broncano, Sarah M. Stark, Caroline M. Schlessman, Michelle R. Raymond, William B. Tran, Adrian D. Kocinski, Nicholas P. Ziats, Wendy A. Goodman

**Affiliations:** Department of Pathology, Case Western Reserve University School of Medicine, Cleveland, OH

## Abstract

Many autoimmune diseases exhibit a female sex bias in prevalence and severity, yet the mechanisms for this remain unclear. 17β-estradiol, a steroid sex hormone with established immunomodulatory roles, signals through the nuclear receptors ERα and ERβ, which are expressed by CD4^+^ T cells. Expression of ERβ is reduced in CD4^+^ T cells isolated from autoimmune disease patients, suggesting that dysregulated E2 signaling contributes to inflammation. We previously identified a novel role for ERβ in promoting the TGFβ-dependent differentiation of Foxp3^+^ Tregs, supporting the idea that ERβ has anti-inflammatory functions. In this study, we investigated the functional role of ERβ in effector T cells, which drive pathogenesis of many autoimmune diseases. We found that CD4+ T cells isolated from mice globally deficient in ERβ exhibit enhanced proliferation and Th1 polarization *ex vivo*, together with elevated levels of proinflammatory cytokines in response to T cell receptor (TCR) stimulation. We also found that transfer of ERβ-KO T cells to immunodeficient mice results in significantly worse inflammation in a murine model of colitis. Together, these findings suggest that T cell-specific ERβ functions as a brake on T cell-mediated inflammation, thereby helping to maintain immune homeostasis.

**Highlights:**

- CD4+ T cells express the nuclear estrogen receptor ERβ
- ERβ-deficient T cells express higher levels of inflammatory cytokines and are hyperproliferative *ex vivo*
- Deletion of ERβ increases T cell-mediated inflammation in a murine model of colitis

**CRediT authorship contribution statement:** SKM: investigation, formal analysis, visualization, writing – original draft; AVB: investigation, formal analysis, visualization, writing – original draft and review and editing; SMS: investigation and formal analysis; CMS: investigation and formal analysis, ADK: investigation, MLR: investigation, WBT: investigation; WAG: conceptualization, funding acquisition, project administration, resources, supervision, formal analysis, visualization, writing-review and editing.

## INTRODUCTION

Nearly 80% of all patients with autoimmune disease are female ^1, 2^, yet the mechanisms underlying this sex bias remain poorly understood. Several factors are known to contribute to sex differences in the immune system, including genetics, environmental and behavioral factors, and the influence of sex hormones ^2, 3^. 17β-estradiol (estrogen, “E2”) is a steroid sex hormone with immunomodulatory functions, independent of its role in female development and reproduction ^4^. Levels of estrogen in females increases significantly during puberty and pregnancy but fall after menopause. These periods of endocrine transition are associated with the remission of the autoimmune diseases rheumatoid arthritis (RA) and multiple sclerosis (MS) ^4^, but the relapse of systemic lupus erythematosus (SLE) ^5^. Similarly, the use of synthetic estrogens, including estrogen-containing contraceptives, is associated with the risk of developing new autoimmune disease, including inflammatory bowel disease (IBD) ^6, 7^. While it is clear that estrogen modulates the immune system, understanding the mechanisms by which estrogen signaling contributes to sexual dimorphism and disease severity in chronic inflammation and autoimmunity is critical to improve patient care and therapeutic approaches.

Estrogen signals through two nuclear receptors, ERα and ERβ, and a membrane receptor GPER1, a G protein-coupled receptor ^8–10^. While estrogen signaling through GPER1 is independent of direct DNA binding and is instead mediated through second messengers, ERα and ERβ are canonical nuclear receptors that mediate estrogen signaling through direct DNA binding ^8–10^. ERα and ERβ bind estrogen in the cytoplasm of a cell, form homo-or heterodimers, then translocate to the nucleus to bind to estrogen response elements (EREs) in the promoters or regulatory regions of target genes ^9, 10^. Estrogen receptors are also able to regulate the expression of genes without an ERE through indirect signaling mechanisms ^8, 10^. In sum, the cellular response to estrogen through ERα and ERβ is dictated by a number of factors, including the expression level of the receptors, homo-vs. heterodimerization, and recruitment of different co-activators and repressors.

Estrogen’s nuclear receptors are expressed across a broad variety of tissues and cell types outside of the reproductive tract, including immune cells ^4, 11^. Our previous work showed reduced expression of ERβ relative to ERα in inflamed mucosal tissues and peripheral CD4^+^ T cells of Crohn’s disease patients ^12^, and similar trends have been observed in other autoimmune diseases including lupus ^13^, suggesting that dysregulated estrogen signaling in CD4^+^ T cells contributes to inflammation. ERα signaling in CD4^+^ T cells promotes activation and proliferation ^14^, similar to its pro-proliferative effect in breast cancer ^15^. Additionally, deletion of ERα has been shown to ameliorate disease in a T cell transfer model of colitis and resulted in decreased serum inflammatory cytokines, worsened histopathology, and decreased the number of IFNγ− and IL-17A-producing CD4^+^ T cells in the lymphoid tissues ^14^. We also previously showed that deletion of ERα was protective in a dextran-sulfate sodium (DSS) model of colitis ^16^, together demonstrating that signaling through ERα promotes intestinal inflammation.

While the function of ERα has been well defined, much less is known about the role of ERβ in CD4^+^ T cells. Deleting ERα limits T cell responses, suggesting that ERβ has a regulatory role. Indeed, our previous study identified a role for estrogen signaling through ERβ in the differentiation and expansion of CD4^+^Foxp3^+^ regulatory T cells (Tregs) *in vivo* and *ex vivo*, supporting a protective regulatory role for ERβ ^12, 17^. In support of this, we also found that Tregs isolated from peripheral blood of Crohn’s disease (CD) patients had reduced expression of ERβ compared to Tregs isolated from healthy controls ^12^. This suggests that loss of ERβ-mediated Treg immunosuppression may contribute to inflammation in CD patients.

In the current study, we asked how ERβ signaling affects the phenotype and function of effector CD4^+^ T cells, and whether this contributes to regulating T cell-mediated inflammation. We hypothesized that ERβ has a critical role in balancing estrogen signaling to maintain immune homeostasis in CD4^+^ T cells, and that loss of this regulation and resultant skewing of signaling through ERα contributes to T cell-mediated inflammation in chronic inflammation and autoimmunity. We first undertook immunophenotyping of mice harboring global deletion of ERβ (ERβ-KO) to gain a broad understanding of the influence of ERβ on the immune system. Deletion of ERβ did not affect the frequency of CD4^+^ T cells or baseline activation of T cells in unchallenged mice. However, ERβ-KO T cells expressed significantly higher levels of proinflammatory cytokines, including MIP-1α, TNFα, GM-CSF, and IL-17, in response to T cell receptor (TCR) stimulation, as well as stronger Th1 polarization and enhanced proliferation in *ex vivo* assays. In line with this, ERβ-KO T cells elicited significantly worse intestinal inflammation and weight loss in a T cell-dependent murine model of colitis, corresponding with accumulation of proinflammatory T cells in the colonic lamina propria of recipient mice. Together, these findings reveal a novel role for T cell-specific ERβ in limiting the pathogenic potential of CD4+ effector T cells, providing protection against chronic intestinal inflammation.

## MATERIALS AND METHODS

### Mice

Wild-type (WT, C57/Bl6 #000664), ERβ-KO (#004745), and RAG2-KO (#008449) mice were acquired from The Jackson Laboratory, bred and housed under specific pathogen free (SPF) conditions, fed standard laboratory chow, and maintained on a 12-hour light/dark cycle. All animal procedures were approved by the Case Western Reserve University Institutional Animal Care and Use Committee (IACUC, protocol #2021-0014).

### T cell isolation

Spleens and mLNs were harvested from WT or ERβ-KO mice, dissociated through a 100 μm strainer, and subjected to red blood cell lysis to prepare single cell suspensions. Cells were counted on a Nexcelom Cellometer using acridine orange and propidium iodide dye (AOPI, Revity). Bulk CD4^+^ T cells were isolated with the Mouse CD4^+^ T cell Isolation Kit (Miltenyi Biotec).

### Immunoblot

Cells were washed once in ice-cold PBS, then lysed in RIPA buffer (Thermo Fisher Scientific) containing protease and phosphatase inhibitors. Protein concentrations were determined using the Pierce BCA Protein Assay (Thermo Fisher Scientific) according to the manufacturer’s instructions. 20-25 μg of protein was loaded on 10% NuPAGE Bis/Tris gels (Thermo Fisher Scientific) for SDS-PAGE. Protein was wet transferred to nitrocellulose and blotted for ERα (Cell Signaling #8644S), ERβ (Invitrogen #PA1-310B), and β-actin (Cell Signaling #12262S). Secondary antibody anti-rabbit-HRP (Cell Signaling #7074S) was used for visualization. Relative protein was quantified by densitometry using ImageJ.

### RNA isolation, reverse transcription, and quantitative PCR

Total RNA was isolated using the High Pure RNA Isolation Kit (Roche) according to the manufacturer’s instructions. For full-thickness colon samples, RNA was isolated using TRIzol reagent (Thermo Fisher Scientific). RNA concentration was measured using a NanoDrop Lite spectrophotometer (Thermo Fisher Scientific). 1 ug of RNA was reverse-transcribed to cDNA using the High-Capacity cDNA Reverse Transcription Kit (Applied Biosystems). Real-time PCR was performed using the TaqMAN Gene Expression Assay with the TaqMAN Fast Advanced Master Mix (Applied Biosystems). Relative gene expression (2^-ddCT^) was calculated using *Gapdh* as a housekeeping gene.

### Flow cytometry

10^6^ cells/sample were washed 1X with PBS, stained with surface antibodies and LIVE/DEAD Fixable Dead Cell Stain (Invitrogen) for 20 minutes at room temperature, and washed with Hank’s balanced salt solution (HBSS) containing 1% FBS before acquisition. For intranuclear staining, cells were then fixed and permeabilized with the TrueNuclear Transcription Factor Buffer Set (Biolegend) or the Phosflow Perm Buffer III (BD Biosciences), then stained with antibodies for nuclear targets. For cytokine staining, cells were re-stimulated with PMA and ionomycin (both Sigma Aldrich) for 5 hours in the presence of Brefeldin A, then fixed and permeabilized with the CytoFast Fix/Perm Buffer St (Biolegend). Samples were acquired using a BD LSR Fortessa and analyzed with FlowJo v10 software (both BD Biosciences).

### Cytokine Array

Bulk CD4^+^ T cells were isolated as described above and stimulated overnight with 3 μg/ml anti-CD3 (Biolegend) and 5 μg/ml anti-CD28 (Biolegend). Supernatants were harvested the next day and clarified by centrifugation. Detection of 40 different chemokines and cytokines was performed using the Proteome Profiler Mouse Cytokine Array Kit (R&D) according to the manufacturer’s instructions.

### T cell Transfer Colitis

As previously described ^18^, CD4^+^CD45RB^high^ cells were FACS-sorted from bulk CD4^+^ T cells isolated from pooled spleen and mLN of WT or ERβ-KO donors. The CD45RB^high^ population was defined as the highest 35% of CD45RB-expressing cells within the CD4^+^ gate. After sorting, cells were washed 1X in cold PBS, counted, and resuspended at a concentration of 4 x 10^6^ cells/ml in PBS. 4 x 10^5^ cells were injected intraperitoneally into RAG2-KO recipients. Recipients were weighed prior to transfer and every other day for 6 weeks. Mice were sacrificed at 6 weeks post-transfer or upon reaching 80% of their initial body weight. Spleen and mLN were collected for cellular analysis by flow cytometry and qPCR as described above. 50% of colons were from each cohort were randomly assigned for histologic assessment and the other 50% were used for isolation of lamina propria mononuclear cells (LPMCs).

### Histological assessment of colonic inflammation

Briefly, colons were flushed to remove feces and filleted longitudinally. Samples for histology were Bouin’s fixed, paraffin embedded, cut to 5 μm, and stained with H&E (AML Laboratories, St Johns County, Florida). Images were obtained using a Zeiss Axiovert microscope and evaluated by a pathologist blinded to mouse genotype for colonic inflammation using an established scoring system ^17^.

### LPMC isolation

To isolate LPMCs, colons were cut into ∼1cm pieces after filleting and transferred to 50 mL conical tubes containing HBSS with 5% FBS and M EDTA. Samples were incubated horizontally at 37C on an orbital shaker at 300 rpm for 15 minutes to remove epithelial cells. Cell suspensions were passed through a 200 μm strainer. The supernatant, containing the epithelial cells, was discarded. The remaining tissue was transferred to a new tube containing fresh HBSS/5% FBS/ M EDTA and incubated for another 15 minutes under the same conditions as before to ensure removal of epithelial cells. The supernatant was again discarded, and the remaining tissue was minced to <1 mm pieces and incubated with Liberase TM (Roche) and DNase1 (Sigma Aldrich) at 37C on an orbital shaker at 300 rpm for 20 minutes. The resulting cell suspension was passed through 100 μm and 40 μm strainers to obtain LPMCs. LPMCs were assessed by flow cytometry and qPCR as described above.

### T cell polarization assays

Naïve CD44^-^CD62L^+^ T cells were isolated from the peripheral lymph nodes of WT or ERβ-KO mice using the Mouse Naïve CD4^+^ T cell isolation kit (Miltenyi Biotec). Cells were cultured with 1 μg/ml anti-CD3 and 2 μg/ml anti-CD28 (both Biolegend) at 1 million cells/ml in RPMI-1640 supplemented with HEPES, sodium pyruvate, L-glutamine, penicillin/streptomycin, and β-mercaptoethanol. To polarize cells, recombinant cytokines were added to the culture as follows: Th1 – rIL-2 (10 ng/mL, Peprotech), rIL-12 (10 ng/ml, Peprotech), anti-IL-4 (10 μg/ml, Biolegend); Th17 – TGFβ (5 ng/ml, R&D), IL-6 (40 ng/ml, Biolegend), IL-23 (10 ng/ml, Biolegend), anti-IFNγ (5 μg/ml, Biolegend), anti-IL-4 (5 μg/ml, Biolegend).

### Proliferation assays

CD4^+^ T cells were isolated as described above and stained with Cell Trace Violet (Invitrogen) according to the manufacturer’s instructions. Cells were cultured for 72 hours with Dynabeads Mouse T-Activator CD3/CD28 (Gibco) at a ratio of 1:2 beads:T cells in the presence or absence of 17β-estradiol (Tocris) at the indicated concentrations. Dye dilution was assessed by flow cytometry after 72 hours on an Attune NXT cytometer (Thermo Fisher Scientific) and analyzed using the proliferation modeling software in FlowJo v10 (BD).

### Data analysis and statistics

Analysis was performed using GraphPad Prism10. Two-tailed Student’s t-test and one-way ANOVA with Tukey’s post-hoc test were used to determine statistical significance. P values <0.05 were considered significant.

## RESULTS

### CD4^+^ T cells express the nuclear estrogen receptors ERα and ERβ

While previous studies have shown that ERα promotes CD4^+^ T cell activation and proliferation ^14, 19^, the role of ERβ in T cells is less understood. Because the cellular response to estrogen is dictated by the level of expression of ERα and ERβ ^8^ [9], we first sought to quantify the expression of ERα and ERβ in primary CD4+ T cells isolated from healthy, C57BL/6 wildtype (WT) mice.

We examined transcript levels of *Esr1* (encoding ERα) and *Esr2* (encoding ERβ) in CD4^+^ T cells from the spleen and mesenteric lymph nodes (mLNs) of male and female mice (10-12 weeks of age). Although *Esr1* expression was similar across all samples (Figure 1A), expression of *Esr2* was elevated in male CD4^+^ T cells from the mLN (Figure 1B). We calculated an *Esr1:Esr2* ratio for each T cell sample, which revealed significantly higher expression of *Esr1* compared to *Esr2* for all samples (Figure 1C). We further assessed protein expression of ERα and ERβ in CD4^+^ T cells from these compartments. In splenic CD4^+^ T cells, we found that ERα expression was more highly expressed in females, but ERβ expression was similar in males and females (Figure 1D-E). Interestingly, mLN CD4^+^ T cells displayed similar ERα expression across males and females, but higher ERβ expression in males (Figure 1F-G). Thus, while ERα may be the predominant nuclear estrogen receptor in CD4^+^ T cells, ERβ is expressed at levels that could influence CD4^+^ T cell phenotype and function.

**Figure 1:**
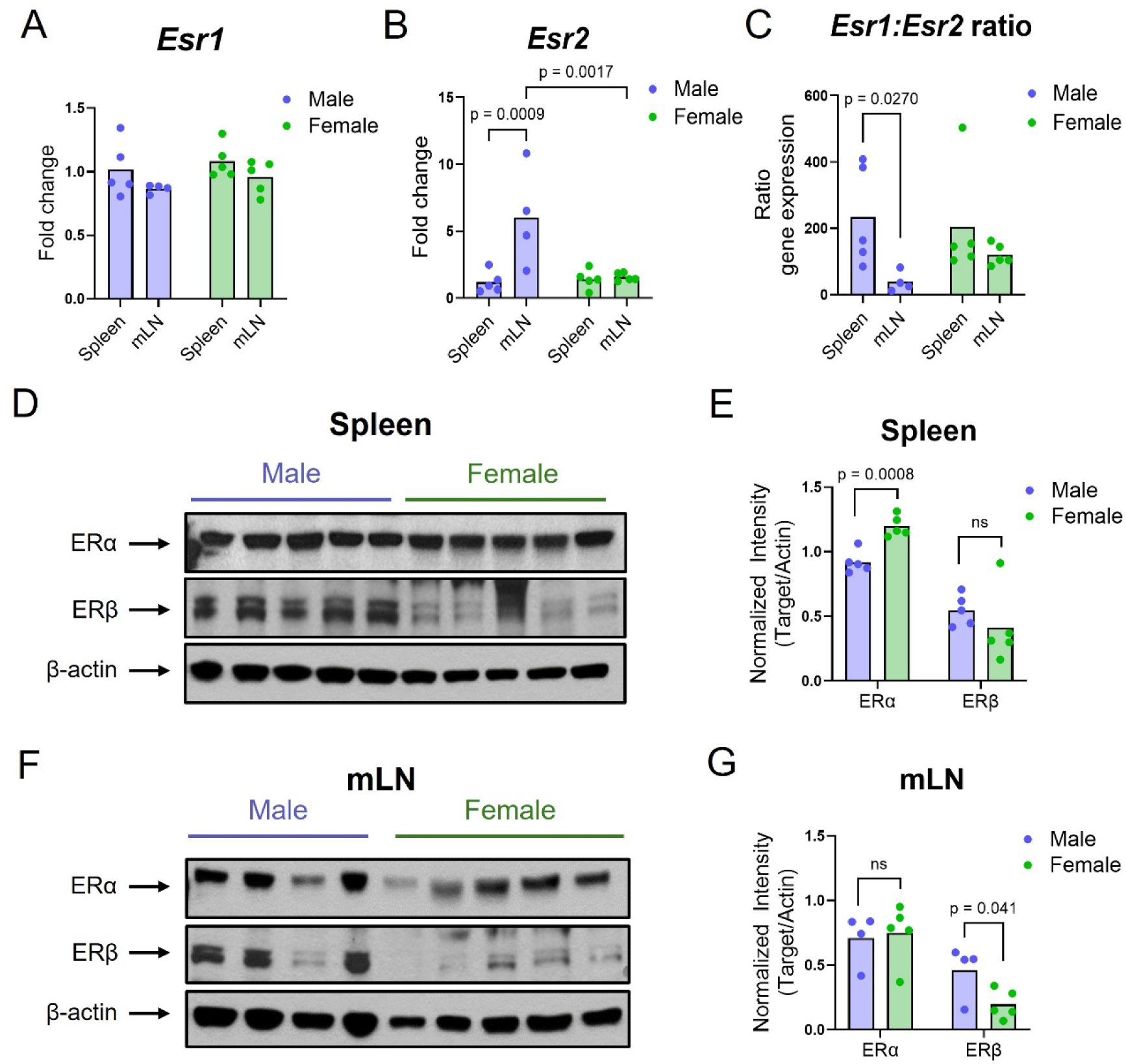
CD4+ T cells express nuclear estrogen receptors ERα and ERβ. CD4^+^ T cells were isolated from spleens or mLNs of indicated mice. Relative gene expression of (A) *Esr1*, encoding ERα, (B) *Esr2*, encoding ERβ, and (C) the ratio of *Esr1:Esr2* were determined by RT-qPCR. Expression of ERα and ERβ were determined by immunoblot of whole-cell protein lysates from (D-E) spleen or (F-G) mLN of indicated mice.

### T cells isolated from unchallenged ERβ-KO mice exhibit normal frequency and activation of peripheral CD4+ T cells

Having confirmed mRNA and protein expression of ERβ in primary CD4+ T cells from WT mice, we next asked if mice lacking global expression of ERβ (ERβ-KO) exhibit changes in T cell frequency or function. First, we assessed expression of T cell populations (bulk CD3+, CD3+CD4+, and CD3+CD8+) in the spleen, mLN, and colonic lamina propria (LPMC) using flow cytometry. WT and ERβ-KO spleen cells displayed similar expression of CD3, CD4, and CD8 in total number and percentage of live cells (Figure 2A-B). Cells from the mLN also had similar expression of these markers, although the total number of CD3-expressing cells was lower in ERβ-KO than WT (Figure 2C-D). LPMCs from WT and ERβ-KO mice also expressed similar proportions of these populations (Figure 2E-F). Altogether, these data show that the frequency of general T cell populations does not differ between WT versus ERβ-KO mice. We further assessed expression of other immune cell types – including B cells, macrophages, monocytes, NK, and NKT cells – in the spleen, mLN, and LPMCs and did not observe major differences between WT and ERβ-KO (Supplemental Figure S1).

**Figure 2:**
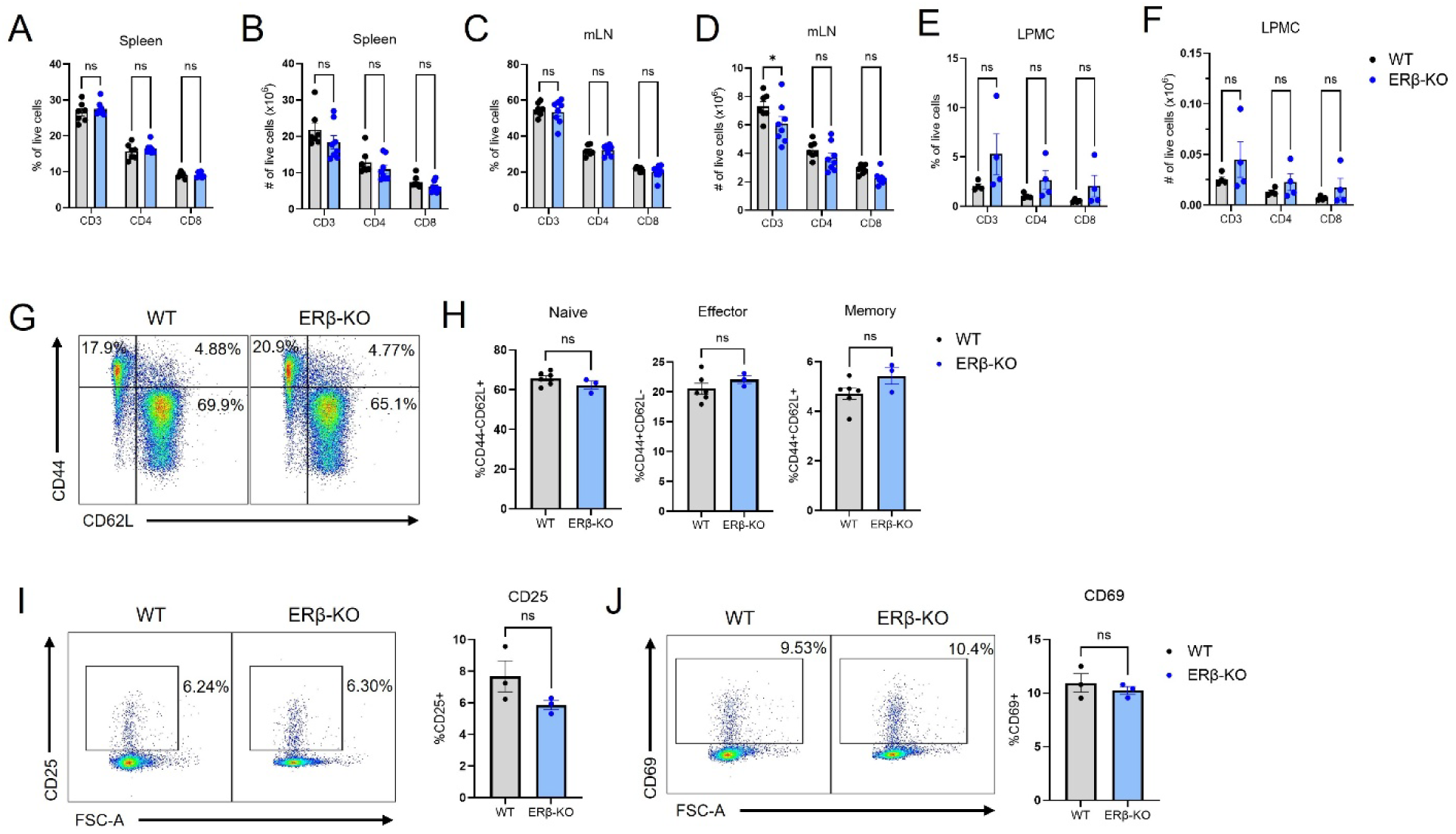
ERβ-KO mice exhibit normal lymphocyte frequency of peripheral T cells. (A-F) Single cell suspensions were prepared from indicated tissues of WT and ERβ-KO mice. Expression of T cell markers CD3, CD4, and CD8 was determined by flow cytometry of spleen (A-B), mLN (C-D), and LPMC (E-F). (G-J) CD4^+^ T cells were isolated from the pooled spleen and mLN of WT or ERβ-KO mice and assessed by flow cytometry for the proportion of (G-H) naïve (CD44^-^CD62L^+^), memory (CD44^+^CD62L^+^), and effector (CD44^+^CD62L^-^) T cells, as well as the expression of activation markers (I) CD25 and (J) CD69. Representative dot plots are shown on the left. Data shown represents mean + SEM, n = 3-8 per group.

ERα has previously been shown to potentiate CD4^+^ T cell activation, with deletion of ERα resulting in a significant reduction in genes, surface markers, and phosphorylation of signaling components indicative of T cell activation ^14^. Given the roles of ERβ in negatively regulating activity of ERα in other cell types, we hypothesized that T cells isolated from ERβ-KO mice may exhibit increases in the proportion of memory/effector cells or higher expression of activation markers. However, we observed similar proportions of naïve T cells (CD4+CD44-CD62L+), effector T cells (CD4+CD44+CD62L-) and memory T cells (CD4+CD44+CD62L+) among WT and ERβ-KO mice (Figure 2G-H). Similarly, there was no significant difference in the expression of activation markers CD25 and CD69 between WT and ERβ-KO (Figure 2I-J).

### ERβ-KO CD4^+^ T cells produce elevated levels of proinflammatory cytokines and are hyperproliferative *ex vivo*

To determine how ERβ influences CD4^+^ T cell cytokine production, naïve T cells (CD4+CD44-CD62L+) were isolated from pooled spleen and mLN of WT and ERβ-KO mice and cultured overnight with anti-CD3 and anti-CD28. Supernatants were collected the next day and assessed for cytokine production using an array with probes for 40 different chemokines and cytokines in technical duplicates. Ten cytokines reached detectable levels in T cell supernatants (Figure 3A), and semi-quantitative analysis of these data using densitometry revealed significance enrichment of MIP1α, TNFα, GM-CSF, IL-17, and IL-3 among ERβ-KO T cells compared to WT, but lower levels of IL-2 in ERβ-KO cells (Figure 3B). Together, these results demonstrate that ERβ limits inflammatory cytokine production in response to T cell receptor (TCR) stimulation.

**Figure 3:**
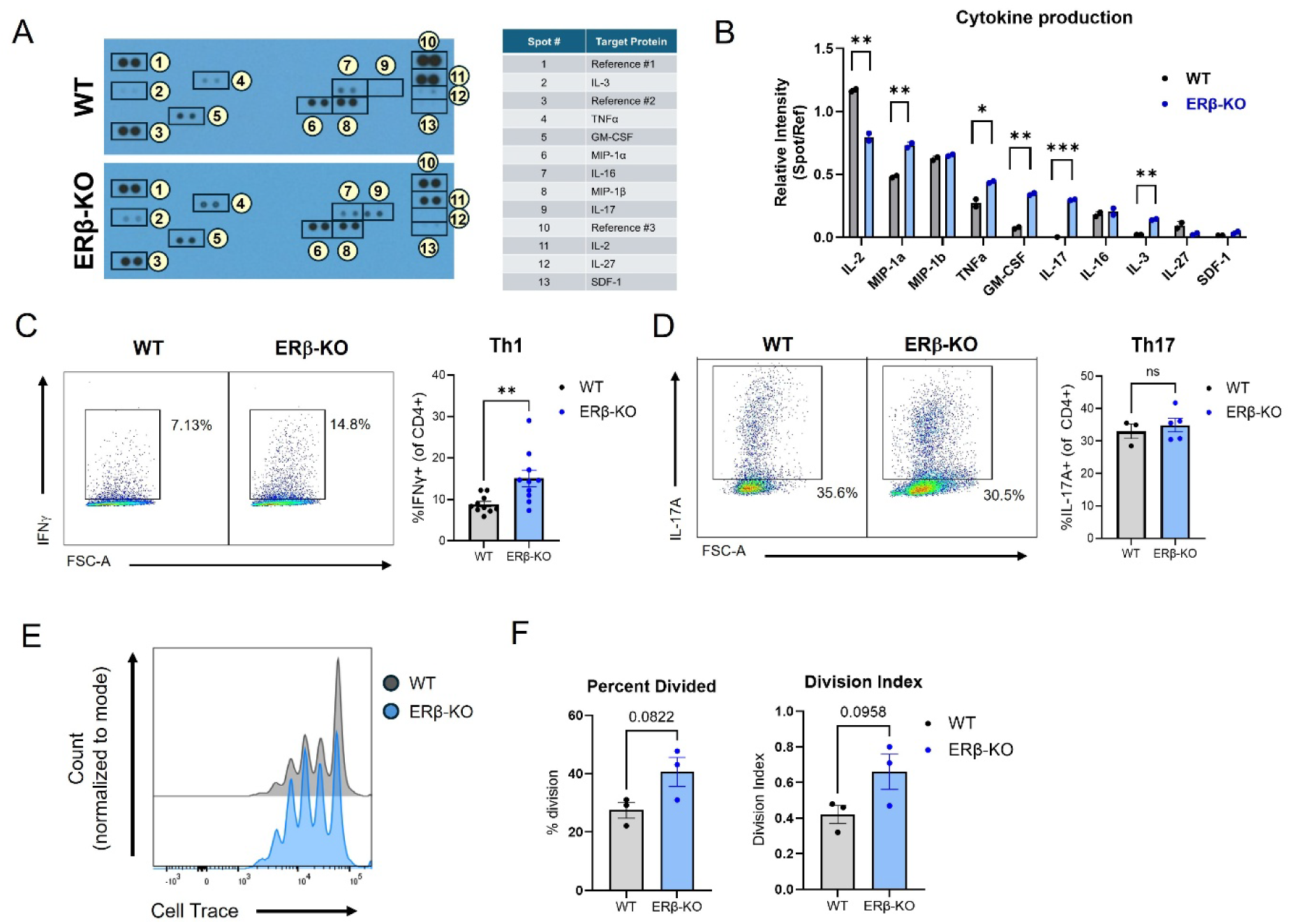
ERβ-KO T cells are hyperproliferative and exhibit altered polarization *ex vivo*. (A-B) WT or ERβ-KO CD4^+^ T cells were stimulated overnight with anti-CD3 and anti-CD28, and (A) supernatants were assayed for 40 different cytokines and chemokines. (B) Relative expression of indicated proteins were quantified by densitometry. (C-D) Naïve (CD44^-^CD62L^+^) CD4 T cells were isolated from the peripheral lymph nodes of WT or ERβ-KO mice and polarized *ex vivo* to (C) Th1 or (D) Th17 lineages. (E-F) CD4+ T cells from indicated mice were labeled with Cell Trace dye and cultured with bead-bound anti-CD3/28 to induce proliferation. (E) Representative histograms show dilution of Cell Trace after 72 hours of culture. (F) Percent of total cells divided and division index are shown. Data shows mean +/− SEM (n=3-10/group). * *p*<0.05, \*\**p*<0.01, \*\*\**p*<0.001.

We next wanted to determine how deletion of ERβ affected CD4^+^ T cell differentiation to proinflammatory Th1 and Th17 populations. To determine baseline *in vivo* differentiation, we collected bulk CD4^+^ cells from pooled spleen and mLN of unchallenged WT and ERβ-KO mice and assessed expression of markers associated with Th1 and Th17 cells. We first analyzed expression of *Tbx21* (encoding Tbet), the master transcription regulator involved in Th1 differentiation, and observed lower RNA expression in ERβ-KO T cells compared to WT (Supplemental Figure S2A). However, the frequency of Tbet+ cells was not different among WT and ERβ-KO by flow cytometry (Supplemental Figure S2B). We similarly assessed expression of the *Rorc* gene and RORγt and found lower gene expression of *Rorc* in ERβ-KO T cells, but no differences in RORγt^+^ T cell frequency by flow cytometry (Supplemental Figure S2C-D). We also analyzed cytokine expression of IFNγ, IL-17A, and IL-10, and observed similar trends of lower RNA expression in ERβ-KO cells but no differences in protein expression (Supplemental Figure S2E-J).

We next tested the potential of naïve T cells to polarize to Th1 and Th17 phenotypes under optimal differentiation conditions, using *ex vivo* Th polarization assays. Naïve CD44^-^CD62L^+^ T cells were isolated from the peripheral lymph nodes (cervical, axillary, brachial, and inguinal) of WT and ERβ-KO mice and cultured with anti-CD3, anti-CD28, and polarizing cytokines to induce differentiation towards Th1 and Th17 lineages. We observed significantly elevated IFNγ production in Th1-polarized ERβ-KO T cells compared to WT (Figure 3C), suggesting that ERβ-KO normally functions to limit Th1 polarization. No differences were observed in IL-17A production among Th17-polarized cells (Figure 3D).

To determine if ERβ has direct role in T cell proliferation, CD4+ T cells were isolated from pooled spleen and mLN, labelled with Cell Trace Violet, and cultured with bead-bound anti-CD3 and anti-CD28 for 72 hours. Dye dilution was assessed by flow cytometry, as shown in representative histograms (Figure 3E). The percentage of cells divided was trending higher in ERβ-KO cells compared to WT (Figure 3F), as was the division index (a measurement of proliferation of the total cell population (Figure 3F). Collectively, these results indicated a potential role for ERβ in limiting T cell proliferation.

### ERβ-KO CD4^+^ T cells exacerbate disease in a murine model of colitis

Based on our findings that ERβ-KO T cells exhibit increased production of classically pro-inflammatory cytokines upon TCR activation, enhanced potential for Th1 polarization, and increased proliferation *ex vivo* (Figure 3), we asked whether these cells may exhibit greater inflammatory function *in vivo*. We utilized the T cell transfer model of colitis, a CD4^+^ T cell-driven model of chronic inflammation in the colon ^18^. As previously described ^20^, CD4^+^CD45RB^high^ cells were FACS-sorted from the pooled spleen and mLN of WT or ERβ-KO donor mice, and 4 x 10^5^ cells were injected i.p into sex-matched Rag2-KO recipients. Recipients were monitored for disease progression and euthanized upon loss of 20% initial weight, or at 8 weeks post-transfer (Figure 4A).

**Figure 4:**
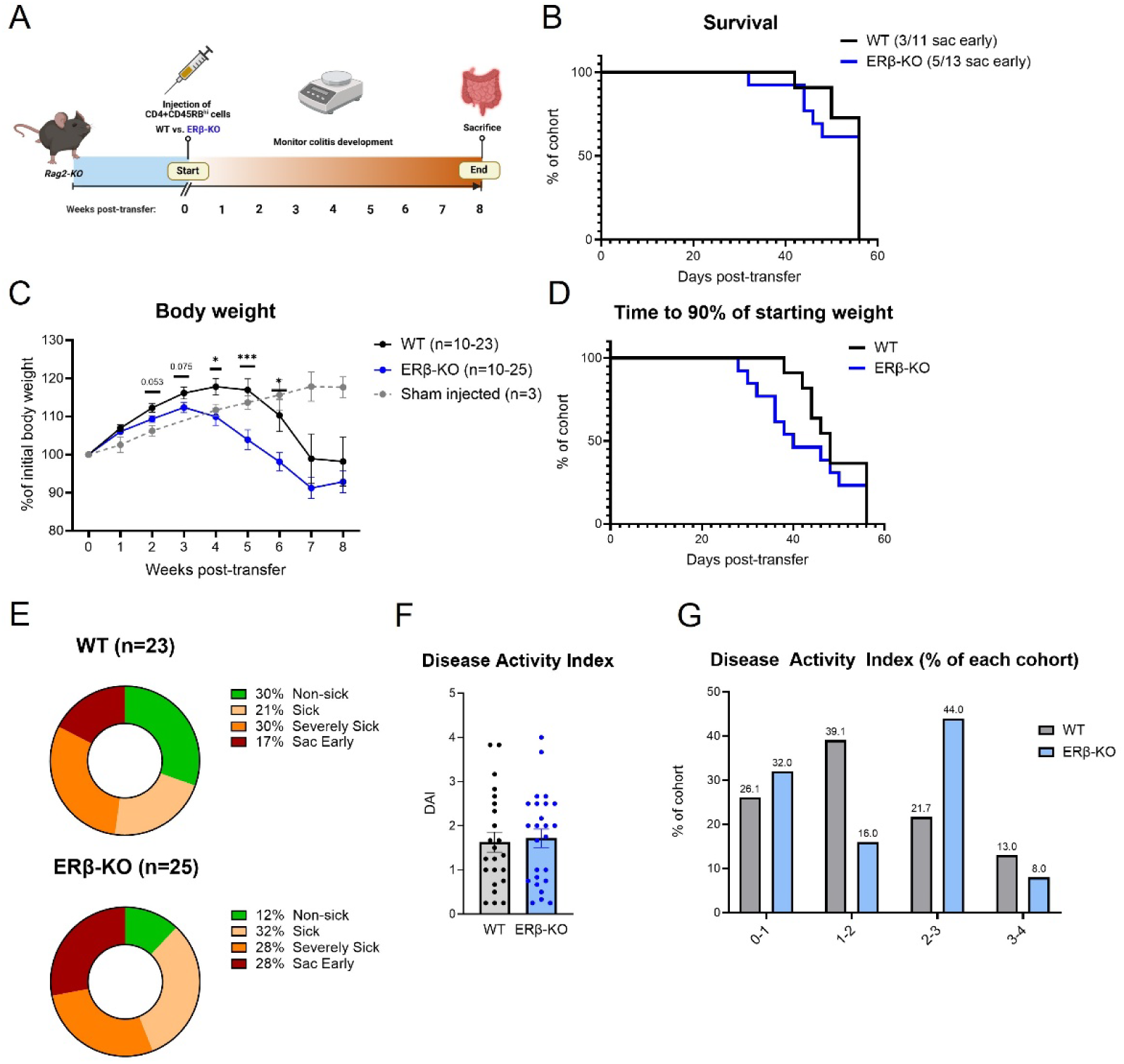
ERβ-KO CD4^+^ T cells confer more severe experimental colitis compared to WT. (A) Schematic of T cell transfer model of colitis. CD4+CD45RB^high^ cells from WT or ERβ-KO donors were adoptively transferred into Rag2-KO recipients and monitored for the development of colitis for up to 8 weeks. (B) Survival curve of the cohorts throughout the time of the study. (C) Weight loss curves showing the changes in body weight after the time of adoptive transfer. Error bars represent SEM. (D) Curve showing the amount of time for recipients to reach 90% of starting weight after adoptive transfer. (E) Disease penetrance measuring the percentage of each cohort that became sick and developed disease. (F) Disease activity index indicating the severity of colitis, consisting of sub scores of the weight loss severity, stool consistency, and presence of fecal blood. (G) The percentage of each cohort within a range of DAI scores. Data shows mean +/− SEM (n=10-25/group). * *p*<0.05, \*\**p*<0.01, \*\*\**p*<0.001.

Although survival was similar across cohorts receiving WT or ERβ-KO T cells (Figure 4B), there were drastic differences in weight loss. Recipients of ERβ-KO donor cells displayed accelerated and exacerbated weight loss compared to recipients of WT cells (Figure 4C), indicating more severe intestinal inflammation in response to transfer of ERβ-KO T cells. This was coupled with recipients of ERβ-KO T cells more rapidly reaching 90% of their starting weight (Figure 4D).

Consistent with this finding, disease penetrance differed among recipients. Recipient mice were categorized as “non-sick” (mice that never lost weight during the 8-week period); “sick” (mice that lost <5% of their initial body weight); “severely sick” (mice that lost between 5-15% of their initial body weight), and “sac early” (mice that lost >15% of their initial body weight and had to be euthanized prior to the end of the study). 30% of mice receiving WT T cells were resistant to colitis development (non-sick), versus 12% of mice receiving ERβ-KO T cells (Figure 4E, green sector). In line with this, 28% of ERβ-KO recipients needed to be sacrificed early, versus only 17% of WT recipients (Figure 4E, red sector).

Disease activity indices (DAI) were calculated for each mouse, comprised of clinical sub-scores measuring the severity of weight loss, stool consistency, and presence of fecal blood. Although the average DAI score was not different between recipients of WT or ERβ-KO cells (Figure 4F), a higher percentage of ERβ-KO recipients had DAI scores of 2 or higher (Figure 4G). Together, these findings suggest that the absence of ERβ confers more severe T-cell driven colitis.

### Colon tissues of ERβ-KO recipients are more inflamed compared to colons of WT recipients

We dissected colon tissues from Rag-KO mice receiving WT versus ERβ-KO T cells 8 weeks post-transfer. Recipients of WT and ERβ-KO T cells displayed similar colon length measurements at the end of the disease course (Figure 5A). Histological evaluation of H&E-stained colon tissues showed that both recipients of WT and ERβ-KO cells exhibited inflammatory damage to the colon, including accumulation of lymphocytes and myeloid cells, neutrophils, goblet cell loss, abnormal crypt architecture, mucosal erosion and ulceration, and transmural inflammation (representative images shown in Figure 5B). Total inflammatory scores (TIS) were calculated as previously described ^17^ and revealed no significant differences between recipients of WT and ERβ-KO cells (Figure 5C and Supplemental Figure S3).

**Figure 5:**
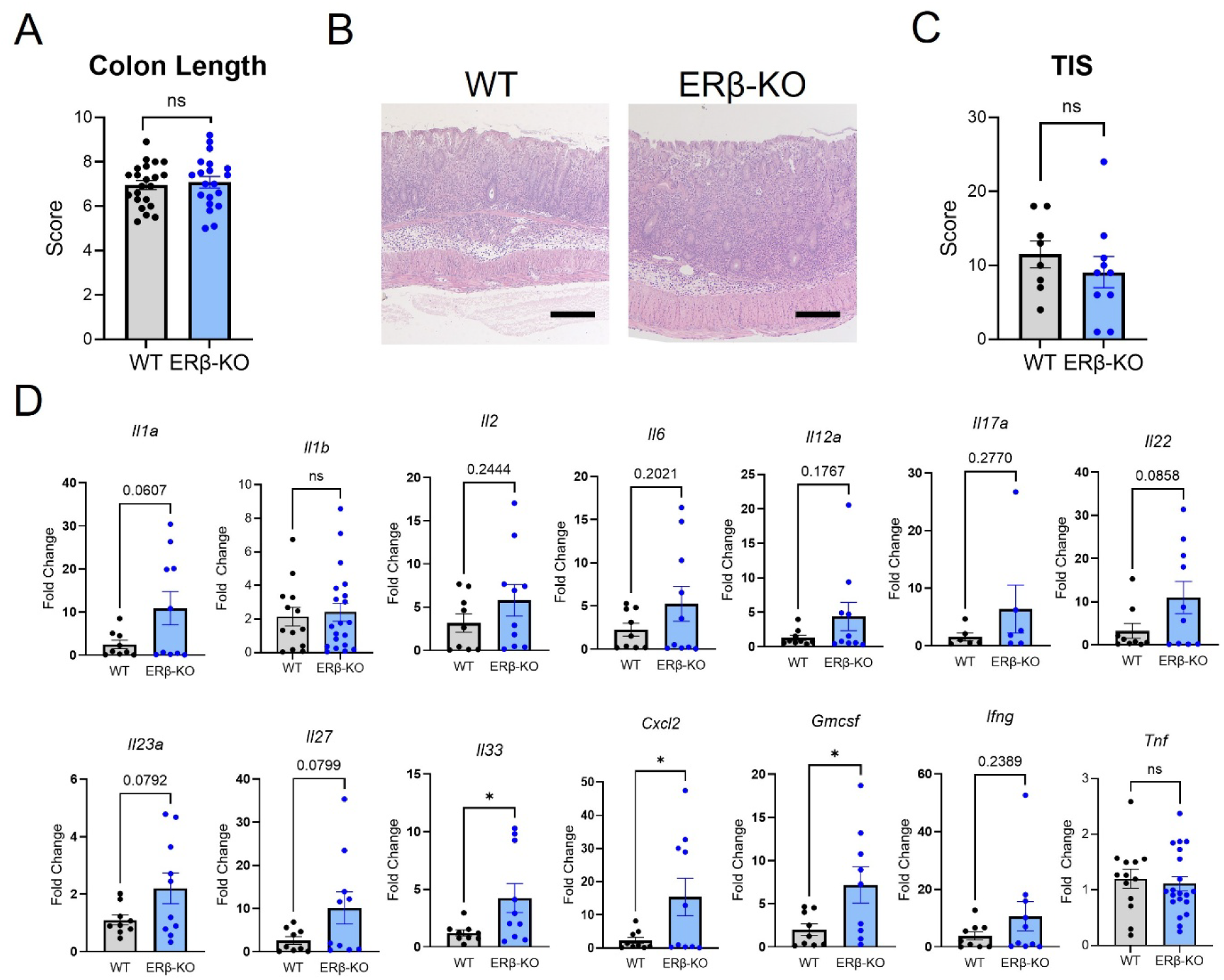
Recipients of ERβ-KO T cells exhibit elevated proinflammatory cytokine production in full-thickness colon tissues. Colon tissues were isolated from indicated T cell transfer mice. (A) Measurement of colon length at the time of sacrifice. (B) Representative images of H&E-stained colon tissues from Rag2-KO recipients at time of sacrifice (scale bar = 100µm). (C) Total inflammatory scores determined by histopathological scoring of H&E-stained sections from recipients of WT and ERβ-KO donor cells. Each dot represents an individual mouse. (D) Gene expression of markers indicating colonic inflammation in full thickness colon tissues taken from recipients of WT or ERβ-KO cells at time of sacrifice. Data shows mean +/− SEM (n=10-25/group). * *p*<0.05, \*\**p*<0.01, \*\*\**p*<0.001.

We also assessed the gene expression of classical proinflammatory markers in full-thickness colon tissues isolated from T cell transfer mice. qPCR analysis revealed that the expression of numerous pro-inflammatory cytokines are elevated in recipients of ERβ-KO cells, with several reaching significance over WT (*Il33*, *Cxcl2*, and *Gmcsf*, Figure 5D). This suggests that T cell-specific ERβ contributes to the regulation of proinflammatory cytokines in the intestine.

### Recipients of ERβ-KO T cells display increased T cell accumulation and activation

As an initial step towards understanding the mechanistic basis for differences in disease severity among our T cell transfer mice (Figures 4-5), we analyzed the CD45RB^high^ T cell populations of WT and ERβ-KO donor mice. Mean fluorescence intensity (MFI) of cells in the CD45RB^high^ gate (top 35% of CD45RB-expressing cells) was comparable between WT and ERβ-KO T cells (Supplemental Figure S4A).

Expression of activation markers CD25 and CD69 (Supplemental Figures S4B-C) and the gut-homing integrin α4β7 (Supplemental Figure S4D) were also similar, indicating that observed differences in response to transfer of WT versus ERβ-KO T cells are not the result of differences in initial expression of CD25, CD69, or α4β7.

We next asked if there are differences in overall cellularity of spleen, mLN, and LPMC among recipients of WT versus ERβ-KO T cells. Tissues were collected and analyzed for total cell counts at the end of the T cell transfer experiment. Interestingly, we observed significantly higher total cell counts in the spleen and LPMCs of recipients of ERβ-KO T cells compared to WT (Figure 6A), suggesting the presence of enhanced inflammation. We also analyzed the frequency of CD4+ T cells in spleens and mLNs of transferred mice, and found increased T cells in spleens of ERβ-KO recipients (Figure 6B). These data indicate that adoptively transferred ERβ-KO T cells have a higher capacity for *in vivo* proliferation.

**Figure 6:**
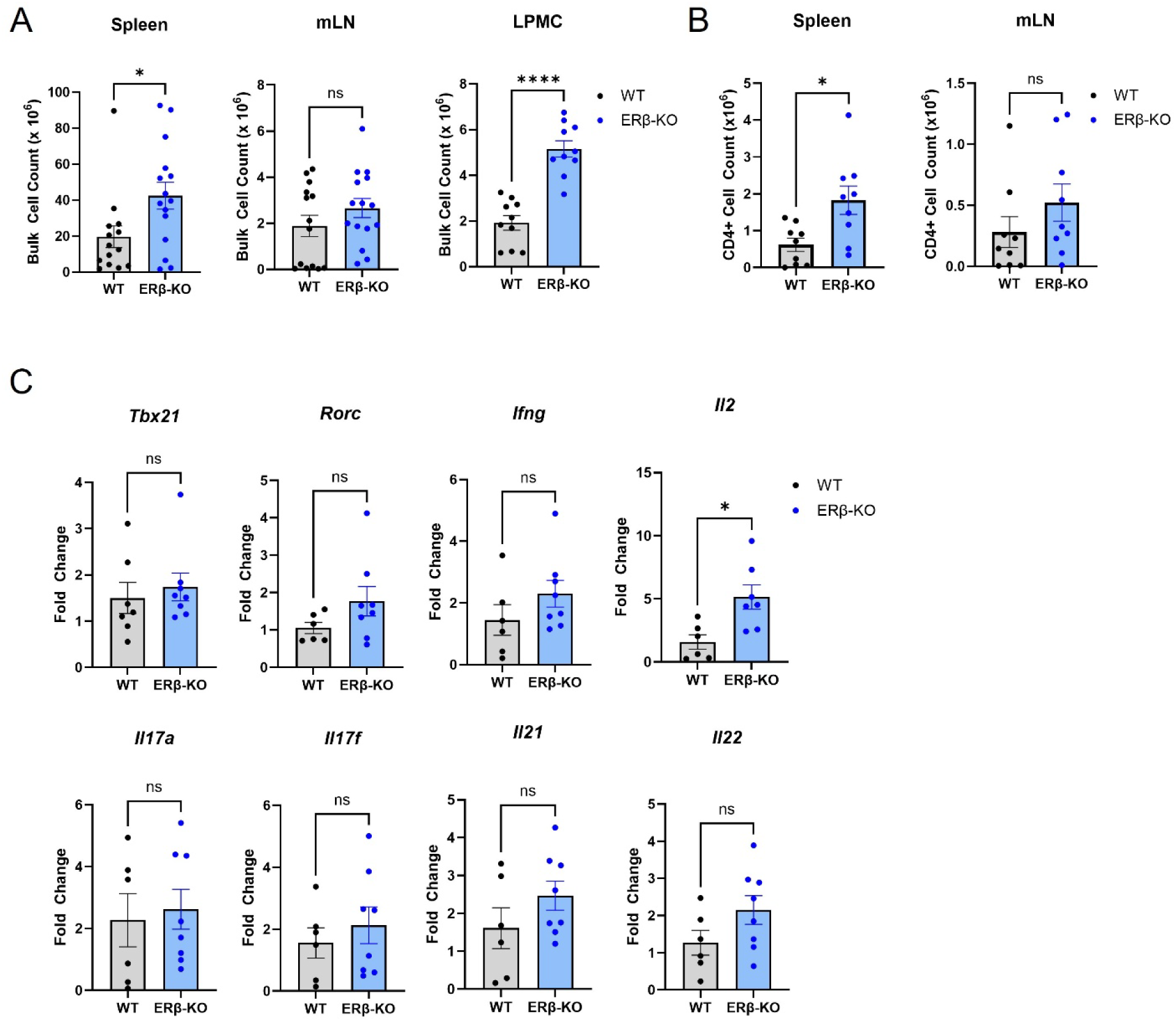
Recipients of ERβ-KO T cells display increased T cell accumulation and activation. (A-C) Total cell counts within indicated tissues at the time of sacrifice. (B) Frequency of CD4+ T cells within indicated tissues at the time of sacrifice. (C) Gene expression of indicated markers among anti-CD3/28-treated T cells isolated from LPMCs of indicated mice at the time of sacrifice. Data shows mean +/− SEM (n=6-15/group). * *p*<0.05, \*\*\*\**p*<0.0001.

Lastly, we isolated CD4+ T cells from colons of transferred mice and rechallenged them with *ex vivo* anti-CD3/CD28 for 16 hours. Gene expression analyses revealed significantly higher expression of *Il2* among T cells isolated from ERβ-KO recipients compared to WT (Figure 6C), along with trends toward higher expression of *Ifng*, *Il17a/f*, *Il21*, and *Il22* (Figure 6C). Collectively, these results suggest that ERβ-KO are pathogenic in vivo via their heightened proliferation and propensity for proinflammatory cytokine production.

## DISCUSSION

The nuclear estrogen receptors have been most heavily studied in the context of breast cancer, where ERα has a strong pro-proliferative role ^15^. Increased interest in the systemic, physiological roles of estrogen has resulted in the identification of ERα and ERβ expression in many tissues and cells outside of the female reproductive tract, including immune cells ^4, 21^. It is not surprising that ERα has a pro-proliferative, pro-inflammatory function in CD4^+^ T cells given its mitogenic role in breast cancer ^14, 19^.

However, much less is known about ERβ in both contexts. It has been suggested that ERβ has unique and opposing functions to those of ERα, suppressing proliferation in breast cancer ^22^. Indeed, we previously identified a key role for ERβ-specific signaling in the TGFβ-induced differentiation of Foxp3^+^ Tregs, supporting this regulatory role of ERβ in the immune system ^12^.

Here, we expanded upon our previous work and showed that ERβ also regulates the phenotype and pathogenic potential of effector CD4^+^ T cells. We hypothesized that ERβ regulates the effect of ERα, which is known to skew CD4^+^ T cells to inflammatory phenotypes, and that deletion of this protective ERβ would exacerbate T cell-mediated inflammation due to enhanced ERα-specific signaling. Our results showed that while ERα is the dominant nuclear estrogen receptor in CD4^+^ T cell, ERβ is also strongly expressed and is capable of influencing T cell responses. However, we did not observe differences in T cell frequency or function in vivo in unchallenged ERβ-KO mice, indicating that in the absence of specific stimuli, ERβ-KO T cells have similar proportions of memory and effector cells and a similar level of activation *ex vivo* compared to WT T cells.

One limitation of the current study was that other cell types that express ERβ may have influenced development or function of the T cells in our global ERβ knockout mice. Publicly available sequencing data (Immgen) has shown that ERβ is expressed in thymocytes prior to the development of mature T cells. Others have identified roles for glucocorticoids and androgens in the thymic epithelial cells (TECs) ^3, 23^, suggesting that estrogen signaling may also influence thymocyte development. Thus, the role of ERβ in T cell precursors and the impact of this signaling on downstream T cell function warrants further study.

Despite negligible differences in activation and proliferation in unchallenged mice, deletion of ERβ increased the differentiation of naïve T cells to IFNγ-producing Th1 cells in our *ex vivo* assays, supporting the idea that signaling through ERβ is anti-inflammatory. Further, ERβ-KO CD4^+^ T cells trended towards increased differentiation of RORγT^+^ Th17 cells and produced more IL-17A and GM-CSF upon activation, suggesting that ERβ has the potential to mediate a variety of inflammatory T cell phenotypes including both Th1 and Th17.

Our *in vivo* model of IBD showed that ERβ-KO T cells induced more severe weight loss and had higher disease penetrance in a T cell-dependent manner, showing that ERβ-mediated regulation is critical for balancing T cell responses during inflammation. These observations are in line with human data showing reduced expression of ERβ in the CD4^+^ T cells of autoimmune patients. However, an important outstanding question is how expression of ERβ becomes reduced in the CD4^+^ T cells of autoimmune patients. One potential answer is the systemic and tissue-level of estrogen, specifically 17β-estradiol (E2), the predominant circulating form of estrogen ^8^. In addition, the other forms of estrogen, collectively termed “pregnancy estrogens”, may also contribute to changes in inflammation observed during pregnancy ^5^. We recently reported that colonic epithelial cells express the necessary enzymes for the conversion of testosterone to estrogen, identifying a previously unknown tissue source of E2 ^24^. The activity of this pathway in the intestine may therefore contribute to estrogen signaling and the downstream T cell phenotype in patients with IBD. Other as-yet unknown tissue producers of estrogen may contribute to other chronic inflammatory diseases.

While many immune genes have EREs and can be directly regulated by ERs ^4^, it is possible that these genes may in turn regulate the expression of the ERs themselves. In breast cancer, members of the forkhead box protein family, including known immune modulators, have been shown to regulate transcription of ERα ^25^. Similarly, Gata3, the transcription factor that drives Th2 differentiation, has also been shown to bind to the *Esr1* gene to upregulate transcription in human breast carcinoma ^25^. Thus, in autoimmune patients, the inflammatory milieu may be responsible for reducing expression of ERβ. Our future studies will seek to address how this inflammation regulates expression of both ERα and ERβ in the context of the immune system.

In conclusion, this study identifies a novel regulatory role for ERβ in the phenotype of CD4^+^ T cells in the context of inflammation, showing that ERβ helps maintain immune homeostasis by limiting inflammatory proliferation and cytokine production. Loss of ERβ expression, as occurs in autoimmune patient T cells ^12, 13^, skews estrogen signaling through the pro-inflammatory ERα, which exacerbates T cell-mediated inflammation ^14^. Future studies to better understand how ERα and ERβ regulate expression and activity of each other will further delineate the roles of these receptors in T cells and other immune cells. Identifying how inflammation may affect expression of ERα and ERβ will also be critical to understanding why expression of ERβ is reduced in the CD4^+^ T cells of patients and how this may be targeted therapeutically to rebalance estrogen signaling and restore immune homeostasis.

## Data availability

All data are available upon request.

## ABBREVIATIONS

CD: Crohn’s disease
E2: estrogen/17β-estradiol
EDTA: ethylenediaminetetraacetic acid
ERα: estrogen receptor alpha
ERβ: estrogen receptor beta
ERE: estrogen response element
FACS: fluorescence-activated cell sorting
FBS: fetal bovine serum
GPER1: g-protein coupled estrogen receptor 1
HBSS: Hank’s balanced salt solution
H&E: hematoxylin and eosin
IBD: inflammatory bowel disease
LPMC: lamina propria mononuclear cell
MFI: mean fluorescence intensity
mLN: mesenteric lymph node
PBS: phosphate buffered saline
SPF: specific pathogen free
TCR: T cell receptor
Treg: regulatory T cell
UC: ulcerative colitis
WT: wild-type

