## Supplemental Figures for "ERβ limits T cell-mediated inflammation to maintain immune homeostasis"

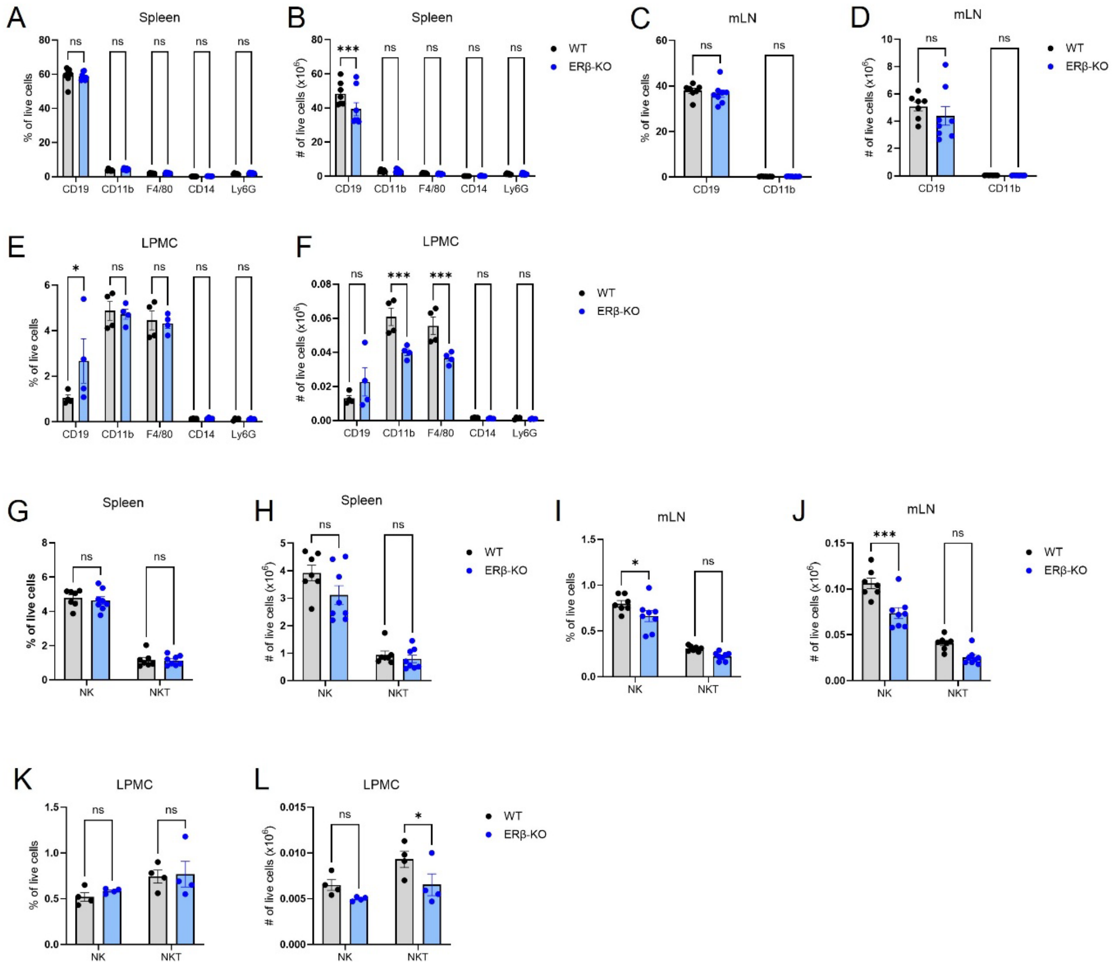


**Supplemental Figure S1: Immunophenotyping of WT and ERβ-KO mice.** (A-F) Frequency and total cell counts of B cells (CD19+), myeloid cells (CD11b+), macrophages (F4/80+), monocytes (CD14+), and neutrophils (Ly6G+) in the (A-B) spleen, (C-D) mLN, and (E-F) LPMC of indicated mice. (G-L) Frequency and total cell counts of NK (CD3-NKp1.1+) and NKT (CD3+NKp1.1+) cells in the (G-H) spleen, (I-J) mLN, and (K-L) LPMCs of WT and ERβ-KO mice were assessed by flow cytometry. Data shows mean +/- SEM (n=10-25/group). * $p$<0.05, **$p$<0.01, ***$p$<0.001.

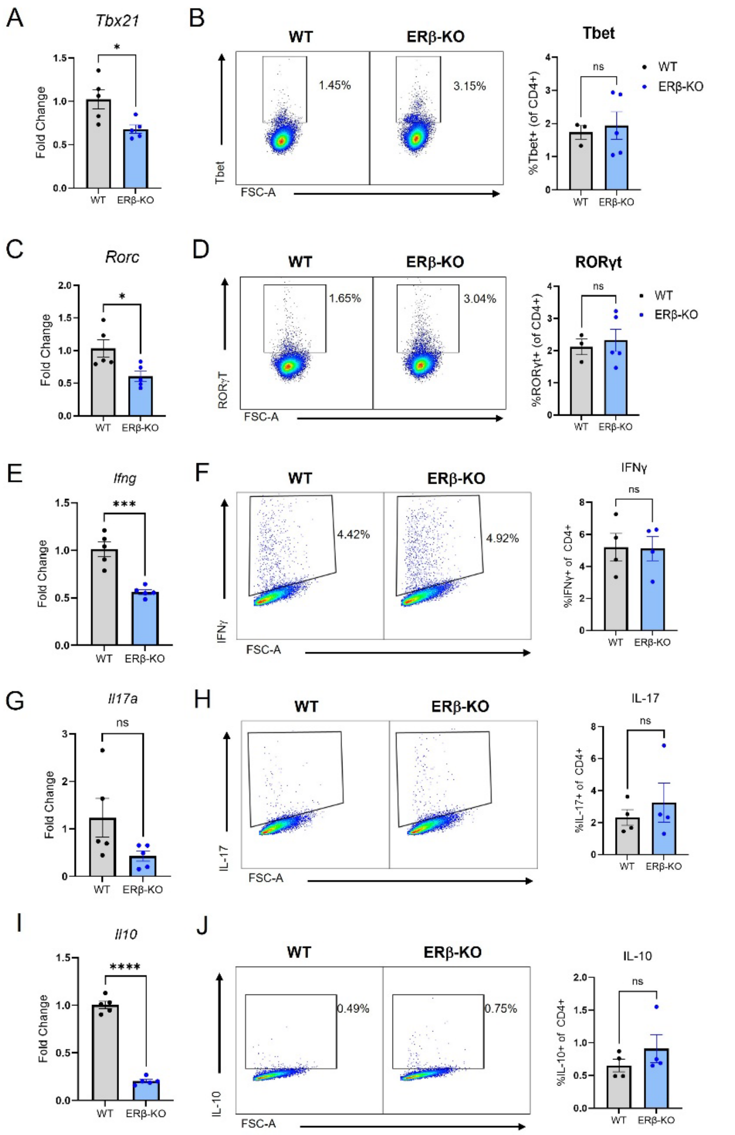


**Supplemental Figure S2: Baseline expression of Th populations in WT and ERβ-KO mice.** Expression of (A-B) Tbx21 and Tbet, (C-D) Rorc and RORgt, (E-F) IFNγ, (G-H) IL-17A, and (I-J) IL-10 was assessed by qPCR and flow cytometry in bulk CD4+ T cells isolated from pooled spleen and mLN of WT or ERβ-KO mice. Data shows mean +/- SEM (n=10-25/group). * $p$<0.05, ***$p$<0.001, ****$p$<0.0001.

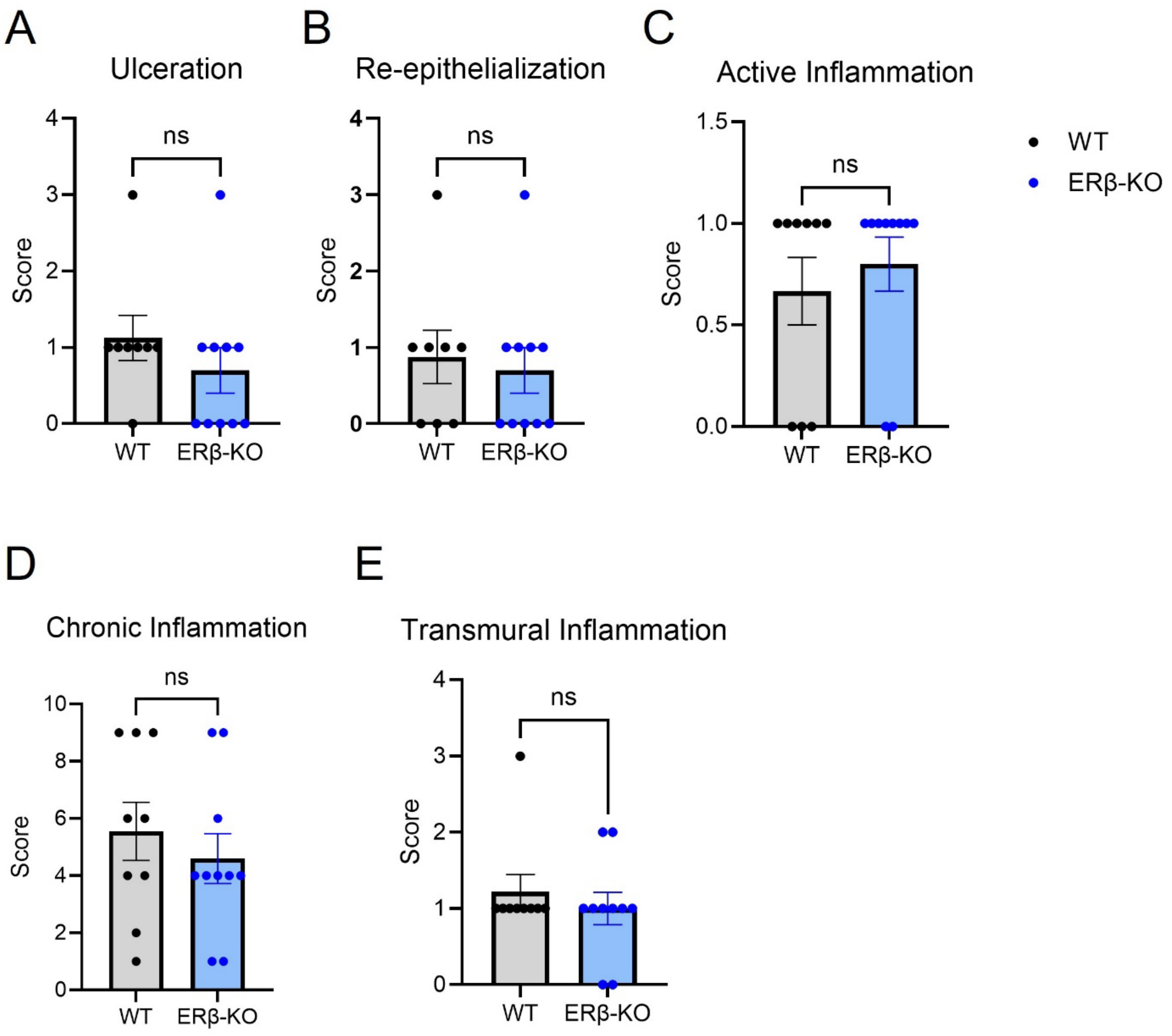


**Supplemental Figure S3: Individual components of Total Inflammatory Scoring for colon histology.** Data shows mean +/- SEM for (A) ulceration, (B) re-epithelization, (C) active inflammation, (D) chronic inflammation, and (E) transmural inflammation in T cell transfer mice. Each dot represents one mouse.

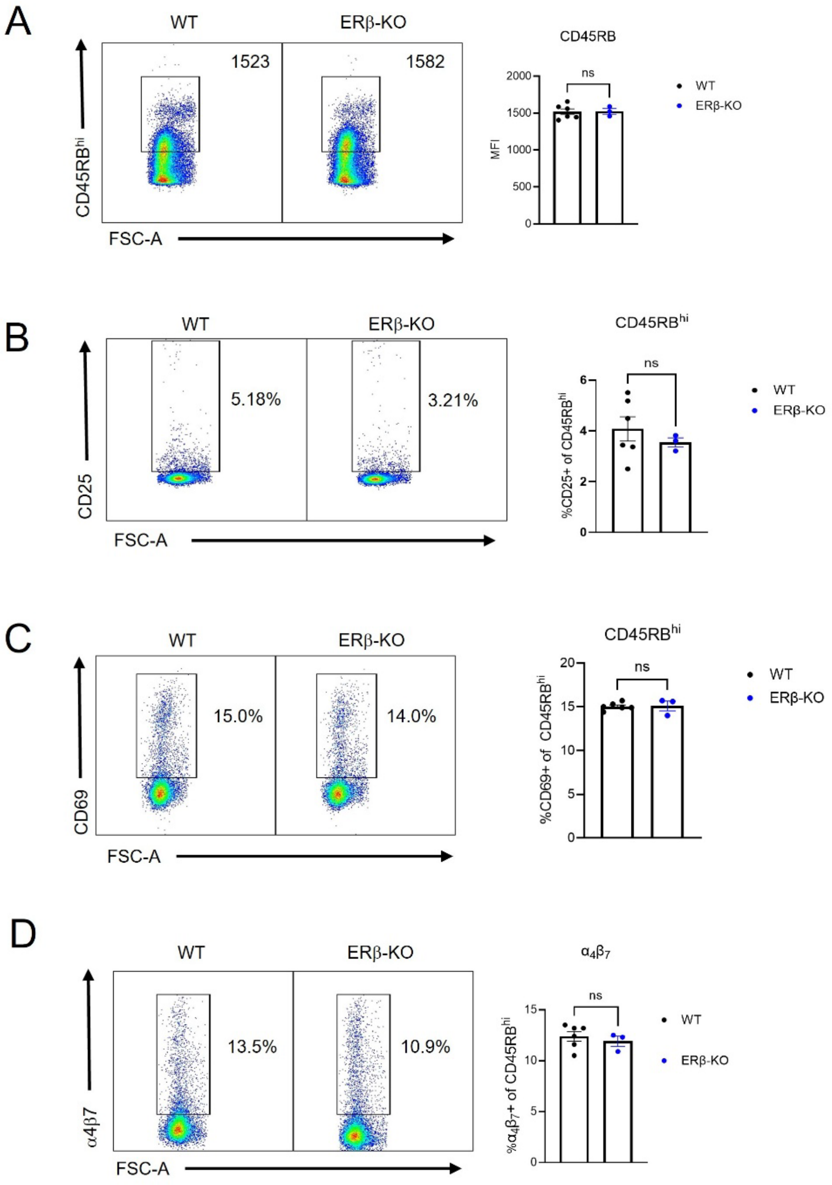


**Supplemental Figure S4: Characterization of CD45RB$^{high}$ cells prior to T cell transfer.** (A) Expression and MFI of CD45RB in bulk CD4+ T cells from pooled spleen and mLN of WT and ERβ-KO mice. (B-D) Expression of the activation markers (B) CD25, (C) CD69, and (D) α4β7 was assessed by flow cytometry. Data shows mean +/- SEM (n=3-5/group).

**Funding:** This work was supported by the National Institutes of Health (R01DK128143 to WAG, T32AI089474 to SKM and AVB); the Crohn's and Colitis Foundation (Senior Research Award #1169305 to WAG); the Cytometry and Imaging Microscopy Shared Resource of the Case Comprehensive Cancer Center (P30CA043703); and the Cleveland Digestive Disease Research Core Center (P30DK097948).

**Data availability:** All data are available upon request.

**Declaration of competing interests:** The authors declare that they have no known competing financial interests or personal relationships that could have appeared to influence the work reported in this paper.
